# Characterization and Mitigation of Fragmentation Enzyme-Induced Dual Stranded Artifacts

**DOI:** 10.1101/2020.01.30.927491

**Authors:** C. Thomas Gregory, Apollinaire Ngankeu, Shelley Orwick, Esko Kautto, Jennifer A. Woyach, John C. Byrd, James S. Blachly

**Affiliations:** Division of Hematology, Ohio State University; Department of Biomedical Informatics, Ohio State University; Department of Pediatrics, Ohio State University; Leukemia Research Program, Ohio State University Comprehensive Cancer Center

**Keywords:** Next-generation sequencing, Enzymatic fragmentation, Sequencing artifacts, Error correction

## Abstract

High-throughput short-read sequencing relies on fragmented DNA for optimal sampling of input nucleic acid. Several vendors now offer proprietary enzyme cocktails as a cheaper and more streamlined method of fragmentation when compared to acoustic shearing. We have discovered that these enzymes induce the formation of library molecules containing regions of nearby DNA from opposite strands. Sequencing reads derived from these molecules can lead to artifact-derived variant calls appearing at variant allele frequencies less than 5%. We present Fragmentation Artifact Detection and Elimination (FADE), software to remove these artifacts from mapped reads and mitigate artifact-related effects on downstream analysis. We find that the artifacts principally affect downstream analyses that are sensitive to a 1-3% artifact bias in the sequencing reads, such as targeted resequencing and rare variant discovery.

**Availability:** Data are deposited at SRA under accession No. PRJNA602687 Software described in this manuscript is freely available at https://github.com/blachlylab/fade

## Background and Introduction

DNA library preparation is a crucial first step in the production of high quality sequencing data. Errors and inefficiencies in the preparation of the library can significantly affect downstream analysis and lead to detrimental effects including increasing the rate of false-positives or false-negatives. Shearing of the physical genetic material is a step shared by most genomic-DNA based sequencing library preparation protocols. This step is performed to provide a uniform DNA fragment distribution, which is ideal for paired-end sequencing.^1^ This shearing is most commonly performed via physical disruption of the DNA by means of focused ultrasonic acoustic waves (“sonication”). Sonication is efficient and consistent, but can be expensive and time consuming. Sonication instruments may cost tens of thousands of dollars and--unless a high-throughput plate-based instrument is employed--samples must be sonicated one to eight at a time for a few minutes each. This time requirement can make large-scale sequencing projects infeasible for small labs, and labor-consuming even for larger operations. Consequently, many groups are exploring alternative methods of DNA fragmentation as a part of the library preparation workflow, including enzymatic fragmentation and transposase-mediated fragmentation-and-tagging (“tagmentation”).

Enzymatic fragmentation employs enzyme cocktails to produce breaks or nicks in the input genetic material. Enzymatic fragmentation can be easily applied to many samples at a time in a 96-well plate based format and does not require specialized equipment.^2^ Enzymatic fragmentation is gaining popularity among high-throughput sequencing operations due to its ease of use, scalability, and low barrier to entry. As enzymes may act non-randomly, however, enzymatic fragmentation could potentially introduce significant biases or sequencing artifacts if the enzymes are blocked from portions of the DNA or selectively shear certain sections of DNA. Current vendors of fragmentation enzyme mixtures for library preparation include Integrated DNA Technologies (IDT; Coralville, IA), KAPA Biosystems (Wilmington, MA), and New England Biolabs (NEB; Ipswich, MA).

KAPA Biosystems markets a proprietary fragmentation enzyme as an alternative to sonication for NGS library preparation as a part of their HyperPlus library preparation kit. This kit effectively combines their popular HyperPrep workflow with a fragmentation enzyme cocktail, the contents of which are proprietary and unpublished. The HyperPlus kit may provide an easier solution to fragmentation than sonication for large cohorts or core facilities. While analyzing sequence variants from a large cohort processed using this kit, we identified a large number of unexpected single nucleotide, insertion, and deletion variants. Upon closer examination of sequence context and alignments within the areas of these false positives, we discovered sequence artifacts that were ultimately a byproduct of the enzymatic fragmentation process. We subsequently tested additional enzymatic fragmentation-based library preparation kits from IDT (Lotus) and NEB, and found similar artifacts, suggesting this is a class effect of the current available commercial enzymatic fragmentation kits. Importantly, these artifacts are also present in public data that utilized these kits^3–7^. We developed a software package, Fragmentation Artifact Detection and Elimination (FADE), to help identify and filter these artifact reads and mitigate their effects on downstream analysis.

## Methods

FADE is written in D, a high-performance statically-typed compiled language, and uses htslib,^8^ the standard library for efficient manipulation of sequencing data files. FADE accepts SAM/BAM/CRAM files containing reads that have been mapped to a reference genome and filters or cleans up artifact-containing reads according to the following procedure. FADE is designed to determine a sequencing read’s enzymatic artifact (EA) status by leveraging aligner soft-clipping. Soft-clipping is an action performed by the aligner to improve the alignment score of a read to the reference by ignoring a portion on one end of the read. Soft-clipping can help an aligner correctly align a read that has sequencing error on one end of the read or has adapter contamination. FADE uses soft-clipping information to identify potentially EA containing reads. First, it will consider only those reads aligned with soft-clipping or containing supplementary alignments; reads with alignments that do not have soft-clipped portions are ignored. Next, a region of the reference sequence of length 600 + L_R_, where L_R_ is the length of the read (not fragment), is extracted such that there exists 300 nucleotides (nt) of padding on each end of the mapped read. Padding of 300 nt on each end of the mapped region provides ample search space for artifact alignment search without being too computationally expensive; most artifacts originate very close to the mapped region and 300 nt was heuristically chosen as an optimal tradeoff, but could be adjusted. The read is next reverse-complemented and then aligned via a Smith-Waterman local alignment^9^ to the extracted region of reference sequence. We use a scoring matrix with a gap open penalty of 10, a gap extension penalty of 2, a mismatch penalty of 3, and a match score of 2. Harsher gap penalties allows the algorithm to be strict in allowing gaps, since we expect the artifact sequences to directly match the reference, except for soft-clipped regions derived from sequencing error. We consider a soft-clipped region to be an artifact if there is a 90% or greater match to the opposite strand sequence.

FADE makes available two modes that rely on the algorithm described above. The **annotate** mode performs the initial analysis and adds BAM tags encoding information concerning artifact status to the alignments, used during filtration to remove the artifacts. The **filter** mode removes reads from the output BAM/SAM file completely if they or their mate contain an identified fragmentation artifact. Optionally, this mode can instead trim artifact-containing reads to remove extraneous sequence (described below), but the reads in total are not removed. After filtration, FADE reports statistics describing the total number of alignments, the percentage of soft-clipped alignments, and the percentage of enzymatic artifacts found. The filtration step must be run on a queryname-sorted BAM file in order to fully filter out the read, its mate, and any other supplementary or secondary alignments.

We used FADE to evaluate chronic lymphocytic leukemia (CLL) and acute myeloid leukemia (AML) patient samples using disease-specific ultra-deep targeted-panel sequencing. CLL samples were prepared using the KAPA HyperPlus kit and KAPA HyperPrep kit with Sonication. AML samples were prepared using the KAPA HyperPlus kit, KAPA HyperPrep kit with Sonication, and the IDT Lotus DNA kit. These samples were sequenced using custom Illumina adapters with dual-barcodes and containing an 8 nt unique molecular identifier (UMI). Reads were trimmed using skewer,^10^ mapped using bwa-mem,^11^ processed according to a modified GATK best-practices pipeline then variant-called using MuTect2.^12,13^ We also used FADE to evaluate sequencing data obtained from the NIH/NCBI Short Read Archive (SRA) from SRA accessions: SRR5009881, SRR5009884, SRR5009885, SRR7665945, SRR7665947, SRR7665951, SRR8695939, SRR8695943, SRR8695947, SRR6389429, SRR6389430, SRR6389431, SRR6911875, SRR6911877, and SRR6911878. These reads were processed using the same pipeline as above. The p-value reported for the 8 CLL samples (Figure 2A) were calculated using a paired t-test. P-values for the SRA samples (Figure 2C) were calculated using pairwise t-tests via ANOVA with p-value correction via the Tukey Honest Significant Difference method^14^.

## Results

Variant analysis on a cohort of cancer samples prepared using the KAPA HyperPlus kit in our lab revealed a large number of unexpected variant predictions that could not be explained initially when viewing the alignments in a genome browser. Upon investigation of these suspect variant calls, we observed that these calls had low variant allele frequencies (VAF) and coincided with subset of alignments in the region bearing soft-clipped ends as shown in Figure 1A. We further determined that these soft-clipped regions were of high base quality, indicating the soft-clipped sequence within these suspect reads were likely derivative of real molecules and not the product of sequencing error. Additionally, the sequences were highly conserved across reads but not representative of adapter sequence. Providing a crucial clue as to their origin, these soft-clipped regions were occasionally identified by the aligner as supplementary alignments of the same reads to the opposite stand. Further investigation revealed that the soft-clipped portion of the suspect reads, if reverse-complemented, could often be found in the nearby reference sequence as shown in Figure 1B.

**Figure 1.**
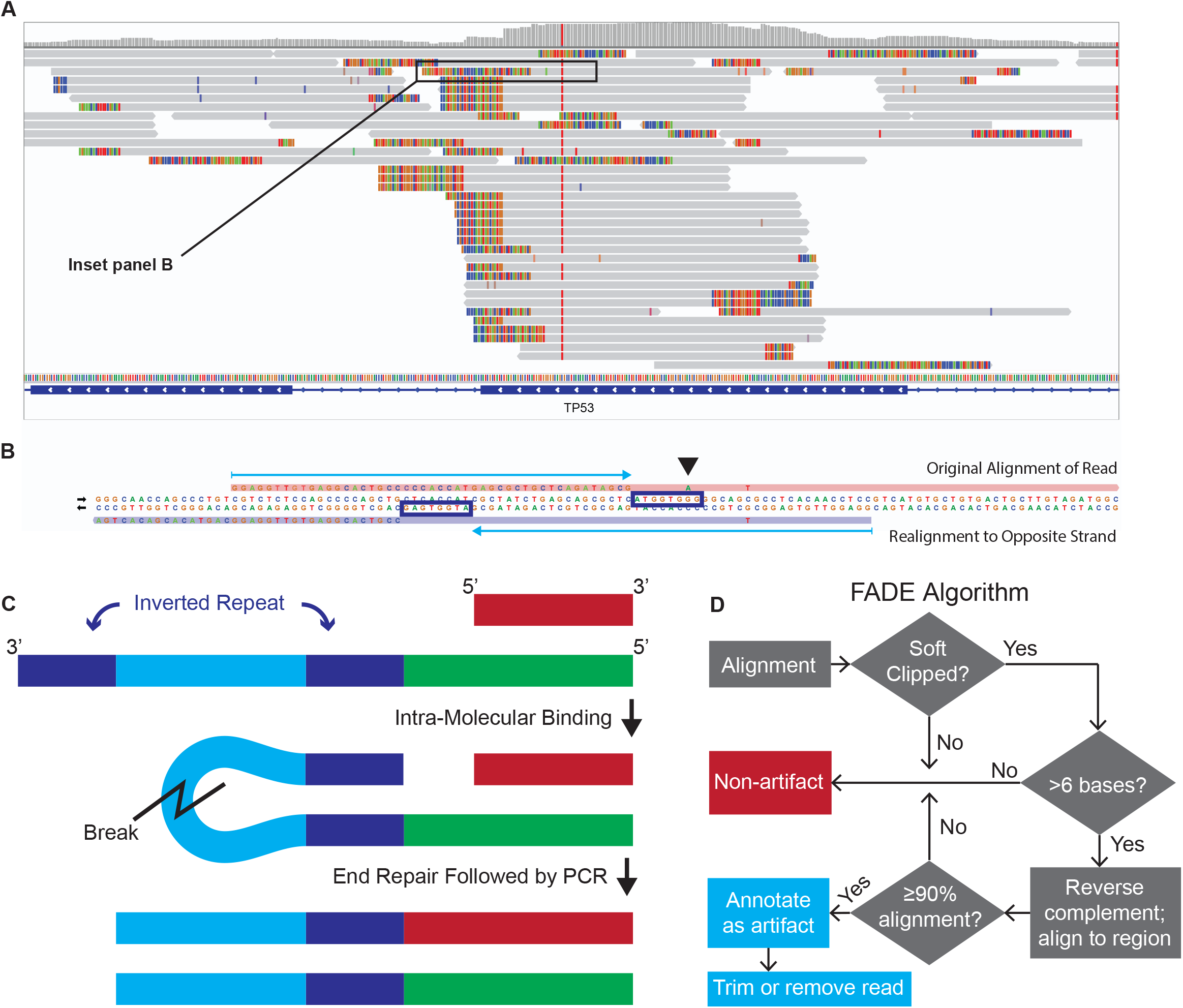
After characterizing the Enzymatic Artifacts (EAs), we hypothesized the mechanism by which they form and developed an algorithm to identify and remove EAs. **A**, Using a genome browser, we observed patterns among reads that contained soft-clipping. Reads without soft-clipped regions are excluded from this view. **B**, Inset from A. After close inspection and realignment of an example read containing a presumed EA, we observed that the soft-clipped region of the read (blue arrow, top) aligned to the opposite strand (blue arrow, bottom). The original alignment and realignment also overlap naturally occurring inverted repeat sequences (purple boxes). Arrowhead: The A mismatch derives from an imperfect repeat and is further demonstration of the mechanistic hypothesis. **C**, We hypothesized that some enzymatic activity exposes these inverted repeats, enabling intra-molecular binding. After end-repair and PCR, this aberration creates DNA molecules that incorporate material from both strands of the original DNA into one strand. **D**, FADE removes enzymatic artifacts by realigning soft-clipped reads to the opposite strand.

Realigning these reads to the strand opposite of their original mapping and in the same region as the original mapping reveals that these artifact reads commonly overlap perfect or near-perfect inverted repeat sequences, which are naturally present in the genome (human in this case, but we have identified this artifact across diverse species). Artifact-containing reads are chimeras: the sequence just proximal to the soft-clipped region aligns with the rest of the mapped strand, while the soft-clipped region of the original read originates from the opposite strand. However, the proximal sequence on the mapped strand is part of the inverted repeat feature from the opposite stand (Figure 1B). A schematic of how this may happen is shown in Figure 1C. Supporting this mechanistic hypothesis, imperfect (e.g, with a single mismatch) inverted repeat features can lead to mismatches from the reference in the non-clipped portion of the read, as the inverted repeats are not a perfect match (demonstrated by the “A” mismatch in Figure 1B). High base quality soft-clipped regions lead INDEL aware variant callers to consider them derivative of a true insertion variant, and explains our high incidence of false-positive INDEL calls (**Suppl. Table 1**).

To identify the cause of these artifacts we permuted several factors in our sequencing workflow, including the individual technician performing library preparation, the specific capture panel, hybridization buffer, thermal cyclers, origin of samples, and sequencing instrument. Most of these factors were ruled out, as we had observed these artifacts in all of our targeted panel-based sequencing across numerous libraries sequenced using a variety of Illumina instruments including Miseq and Hiseq 4000. With all other variables in the workflow addressed, we turned our attention to the sequencing library preparation kit. Focusing on the enzymatic fragmentation aspect of library preparation, we prepared libraries from the same samples, but using Covaris sonication in place of the enzymatic fragmentation.

FADE analysis on matched samples prepared with both KAPA HyperPlus (enzymatic fragmentation) and KAPA HyperPrep (sonication) found a clear difference in the number of reads identified as containing the described artifact (Figure 2A). The artifact rate, defined as the percentage of all mapped sequencing reads identified as containing artifact, was about 2% in samples sequenced using Kapa HyperPlus, whereas the artifact rate of samples subjected to sonication was about 0.01%. Analysis of a 1300 sample patient cohort with FADE revealed artifact in all samples sequenced using the Kapa HyperPlus kit at a level of one to three percent (Figure 2D). The 0.01% artifact rate in the sonicated samples is likely false positive for reads originating from repetitive regions of the genome or misalignment of the Smith-Waterman alignment that still passes the scoring threshold.

**Figure 2.**
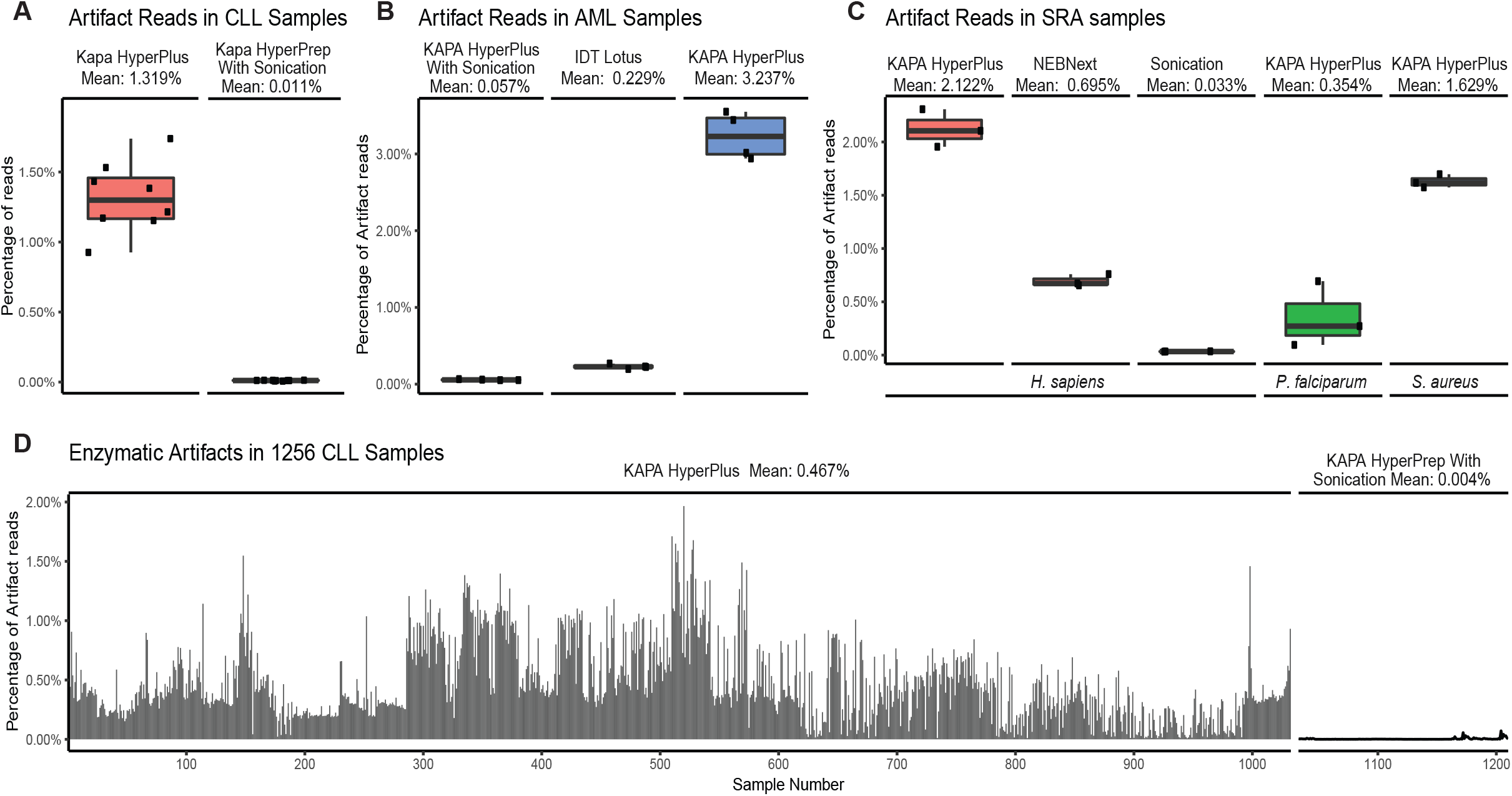
FADE was used to demonstrate the effects of enzymatic fragmentation on a variety of samples and conditions. **A**, After running FADE on targeted-capture samples prepared using both the KAPA HyperPlus kit and the KAPA HyperPrep kit with sonication, we found an average 1.3% difference (p <0.001) in artifact rate reported by FADE between the HyperPlus and the sonication groups. **B**, AML samples were also tested using sonication, the KAPA HyperPlus kit, and the IDT Lotus kit. **C**, FADE analysis of samples from SRA. Depicted are 3 *H. sapiens* (WGS, KAPA HyperPlus), 3 *H. sapiens* (WES, NEBNext), and 3 *H. sapiens* (WGS, Sonication) samples; as well as 3 *P. falciparum* (WGS, KAPA HyperPlus) and 3 *S. aureus* (WGS, KAPA HyperPlus) samples. With the enzymatic fragmentation samples, we observed an artifact rate of 0.2 to 2.5 percent. Human samples using KAPA HyperPlus had an average difference in artifact rate of 2.08% when compared to the human sonication samples. All enzymatic fragmentation artifact rates from the human samples were significantly higher than the sonicated artifact rates (p <0.0001). **D**, Among our own large CLL targeted re-sequencing cohort, we observed a similar improvement when changing our workflow from enzymatic fragmentation to sonication.

This discovery implicates the fragmentation enzyme in the creation of dual stranded library molecules that do not represent the native DNA state. Reads containing this extraneous sequence typically originate from region of the genome with inverted repeat sequences. We hypothesized that enzymatic fragmentation cocktails occasionally generate large (10-30 nt) sticky ends that expose these inverted repeat sequences inducing the formation of a stem-loop structure that persists into amplification (illustrated in Figure 1C). Leveraging our characterization of the artifact, we created FADE to identify artifact-containing reads and remove or trim those reads; the process is outlined in Figure 1D. With FADE, we were able to identify the origins of the soft-clipped molecules and exclude these reads from downstream analysis, or remove only the EA-containing soft-clipped portion.

To identify whether this issue is isolated to one particular product, we next tested IDT’s Lotus DNA kit on our AML test samples. As shown in Figure 2B, the Lotus kit appears to have an artifact rate higher than that of sonication, however the HyperPlus kit has a relatively worse artifact rate than that of Lotus. This indicates that the issue is not limited to a specific manufacturer’s enzymatic fragmentation process, and is likely characteristic of the class of enzymes being used across the field.

To test the hypothesis that these artifacts were not a result of some process local to our operational procedures, we analyzed Next Generation Sequencing (NGS) data from projects on the Sequence Read Archive (SRA) that were reported to have been prepared using enzymatic fragmentation or sonication. To further test whether the artifact may be a byproduct of hybrid capture targeted resequencing, we selected projects that were whole-genome sequencing (WGS) or whole-exome sequencing (WES) based. For each of three samples from each project, we performed adapter trimming, filtering of low-quality reads, mapping to a reference genome, and then analyzed them for artifacts using FADE. As shown in Figure 2C, FADE detected double-stranded DNA library artifacts in all tested sequencing experiments from the Sequence Read Archive that used the KAPA enzymatic fragmentation at levels comparable to what has been witnessed in our larger cohorts. In contrast, randomly selected sonication samples from SRA had an artifact rate of about 0.01%, whereas enzymatic fragmentation samples varied from one to three percent (Figure 2C). Additionally, we observed that SRA samples prepared using NEB’s NEBNext enzymatic fragmentation kit were found to have artifact rates greater than that of sonication but less than that of the HyperPlus kit, at 0.69%. At the time of manuscript preparation, we were unable to locate samples in SRA that stated use of the IDT Lotus kit.

As the ultimate goal of many DNA sequencing experiments is the detection of DNA variation, including somatic mutations, we next examined variant calls across a panel of library preparations. Figure 3A shows the resulting false positive variant calls incurred by enzymatic fragmentation in these samples. Using FADE to identify and remove artifact-containing reads from the AML samples, we compared variant calls before and after using FADE. We found that up to 60% of variant calls could potentially be attributed to EA-containing reads. The bulk of these variants, however, fall below 5 percent VAF as shown in Figure 3B.

**Figure 3.**
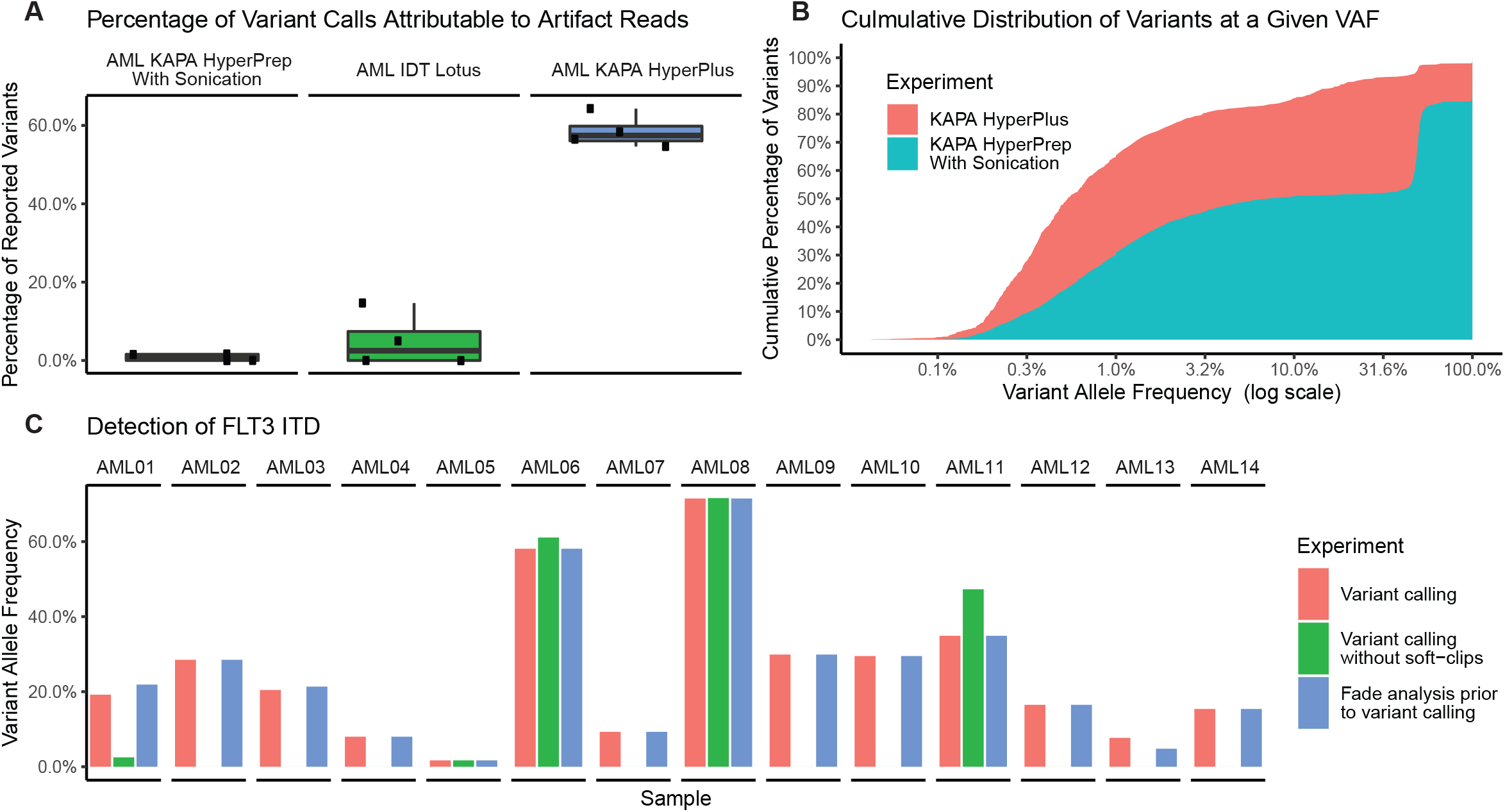
Though only a small percentage of reads are affected by enzymatic fragmentation, the effects on variant calling can be significant. **A**, The samples from **Figure 2b** underwent variant analysis with MuTect2. Here, we show the percentage of variant calls that went unreported after performing artifact removal with FADE. **B**, Cumulative variant allele frequency distributions are shown for sonication and enzymatic fragmentation samples from a cohort of targeted resequencing samples. Enzymatic fragmentation samples show a much larger distribution of variants below 10% VAF than the sonication samples. **C**, Using MuTect2, we detect the *FLT3*-ITD in all 14 AML samples. Ignoring soft-clipped regions removes the erroneous effects of enzymatic fragmentation, but results in lost detection of the ITD in 9 samples. FADE removes these biases while retaining the power to detect the *FLT3*-ITD in all cases.

Because clinically actionable variants may be present at low VAF in contexts including drug resistance^15^ and measurable residual disease,^16^ a hard floor cutoff for VAF may be inappropriate. We next looked to see whether unique molecular identifier (UMI) consensus calling would assist in the calling of low VAF mutations while avoiding false-positive variants originating from artifact containing reads. We observed that UMI consensus calling did not substantially influence the artifact rate **(Suppl. Figure 1).** This result supports the hypothesis regarding EA generation, as fragmentation (with presumptive loop formation and breakage) occurs before UMI-containing adaptors are ligated to the library. Thus, any artifact originating from physical DNA structure created prior should persist through NGS library preparation. We conclude that using UMIs for error correction and variant resolution below 1% variant allele frequency in conjunction with enzymatic fragmentation may be misleading, as the described artifacts would still contaminate true variant results.

To address the concern that EAs could be mitigated by simply ignoring reads containing soft-clipped regions during variant calling, we considered small to intermediate structural variant calling, which may rely on the presence of soft-clipped regions. Internal Tandem Duplication (ITD) in the *FLT3* gene (*FLT3*-ITD) is of high clinical relevance in acute myeloid leukemia^17^ and is detectable even with short-read sequencing. Because MuTecT2 and other popular variant callers use soft-clipped regions to help identify insertion variants, we tested the effect of using the `dontUseSoftClippedBases` flag on the detection of the ITD in AML cases confirmed positive for *FLT3*-ITD and that are detectable using MuTect2 with default settings. In 9 of 12 cases MuTect2^12^ was unable call the *FLT3*-ITD when ignoring soft-clips as shown in Figure 3C. To ensure that FADE did not indiscriminately remove true-positive structural variants (*FLT3*-ITD), we performed FADE analysis and removal followed by variant calling. After removing EAs followed by MuTect2’s default variant calling, MuTect2 retained the ability to call *FLT3*-ITD variants in all 12 cases.

## Discussion

Enzymatic fragmentation of DNA offers a cost-effective and simple alternative to acoustic shearing. This is attractive for both resource-limited settings as well as large, high-throughput operations. Further, focused ultrasonic waves have even been shown to induce DNA damage (8-oxoguanine) that results in a misread base on modern sequencing instruments;^18^ some software now considers this phenomenon when issuing variant calls.

Here, we discovered a surprising number of reads from high-depth, targeted capture experiments contained sequence from both strands in a consistent and predictable pattern. These DNA sequencing libraries have been prepared by different individuals, on different targeted re-sequencing panels, using different capture buffers, using many different thermal cyclers, and have originated and been isolated from a variety of different research lab environments and research groups. Through a process of elimination, we isolated the origin of these reads to library preparations that included an enzymatic fragmentation step.

Due to the proprietary nature of the constituents comprising the enzymatic fragmentation cocktails in the kits we tested, it is difficult to provide a mechanism for the formation of the artifacts or solution to eliminate them. Because of the effects on variant predictions, the high base quality of these artifacts, its presence in all of our sequencing data using enzymatic fragmentation kits, its presence in public datasets that use enzymatic fragmentation, and the change in artifact rates when using sonication on matched samples, we conclude that NGS library molecules are produced that are not representative of the original DNA and are a byproduct of enzymatic fragmentation. The percentage of artifactual reads created by the enzymatic fragmentation workflow is about one to three percent or less of the total reads.

Plate-based enzymatic fragmentation is more cost effective and less time-intensive than sonication. Unfortunately as we have shown here, it lends itself to low percentage biases. Sonicated DNA is also prone to oxidative DNA damage yielding false positive variant calls, however, some variant callers provide algorithms to tag potentially suspect variant calls resulting from this damage.^19^ For other purposes, e.g. WGS or WES without rare variant analysis, and other experiments that are less sensitive to a roughly one to three percent bias, enzymatic fragmentation kits streamline NGS library prep in a time and cost-effective manner. In WGS, the artifact-containing reads are dispersed across the genome, compared to targeted panel sequencing where artifact-containing reads may accumulate at fewer loci and more adversely affect variant calling. In targeted panel sequencing, a specific artifact location is more likely to be sampled multiple times during library preparation and more likely to show up in downstream analysis. It is likely EAs as described here have gone unnoticed due to their minimal effect on WGS and WES data which are not generally used for rare variant analysis.

One cannot simply remove EA-derived variants based on VAF, as there exist clinically actionable variants at VAFs comparable to the frequency of EAs. UMI consensus calling can help resolve down to 0.01 VAF with confidence by polishing out sequencing error,^20^ but EAs originate from a physical molecule prior to the ligation of UMIs and thus this strategy is also unhelpful in the elimination of EAs. INDEL-aware variant calls may produce extraneous variant calls by considering the soft-clipped sequences as true INDEL variants. Imperfect inverted repeat sequences (i.e., containing mismatches, Figure 1B) may also yield false-positive variant calls. However, it is not feasible to ignore soft-clipped reads when variant calling to avoid artifact reads. INDEL-aware variant callers such as MuTect2 use soft-clipped reads to help identify true INDEL variants, like the clinically relevant *FLT3*-ITD.^17^ We showed here that when ignoring soft clipped bases, we are unable in most cases to detect this structural variant in samples with a confirmed and otherwise-detectable *FLT3*-ITD. FADE removes artifact reads but still allows detection of *FLT3*-ITD; this is likely similar for other intermediate insertion-type variants and other SVs.

As the EA-derived variant calls tend to occur at VAF less than 5%, we recommend performing FADE analysis to remove artifact reads when rare variants are of interest. For those that wish to use enzymatic fragmentation kits in conjunction with rare variant analysis or reanalyze existing datasets created with enzymatic fragmentation, we make FADE freely available on GitHub at https://github.com/blachlylab/fade.

## Acknowledgements

We are grateful to the patients who graciously donated tissue samples for research.

This work was supported in part by NIH 5P30CA016058, NIH R35CA197734, NIH R01CA183444, and a grant of equipment from StorageReview.

## Contributions

T.G. and J.S.B. conceived and designed the study. J.A.W, J.C.B., and J.S.B. contributed essential study reagents and patient samples. T.G., A.N., and S.O. contributed to methodology and experimental design and performed research. T.G. and E.K. developed algorithmic methods and contributed to data analysis and curation. J.S.B provided project supervision and project administration. All authors contributed to and edited the manuscript, and approved the final version of the manuscript.

