## Supplementary Data for "Characterization and Mitigation of Fragmentation Enzyme-Induced Dual Stranded Artifacts"

| sample | INDEL RATE | SNP RATE | INDEL Number | SNP Number | Fragmentation |
| --- | --- | --- | --- | --- | --- |
| AML_1a | 0.183673 | 0.816327 | 9 | 40 | Sonication |
| AML_2a | 0.164179 | 0.835821 | 11 | 56 | Sonication |
| AML_3a | 0.265625 | 0.734375 | 17 | 47 | Sonication |
| AML_4a | 0.133333 | 0.866667 | 8 | 52 | Sonication |
| AML_1b | 0.589286 | 0.410714 | 66 | 46 | KAPA Hyper Plus |
| AML_2b | 0.576087 | 0.423913 | 53 | 39 | KAPA Hyper Plus |
| AML_3b | 0.488095 | 0.511905 | 41 | 43 | KAPA Hyper Plus |
| AML_4b | 0.490741 | 0.509259 | 53 | 55 | KAPA Hyper Plus |

Supplementary Table 1: INDEL and SNP rates among sonication and enzymatic fragmentation processed samples.

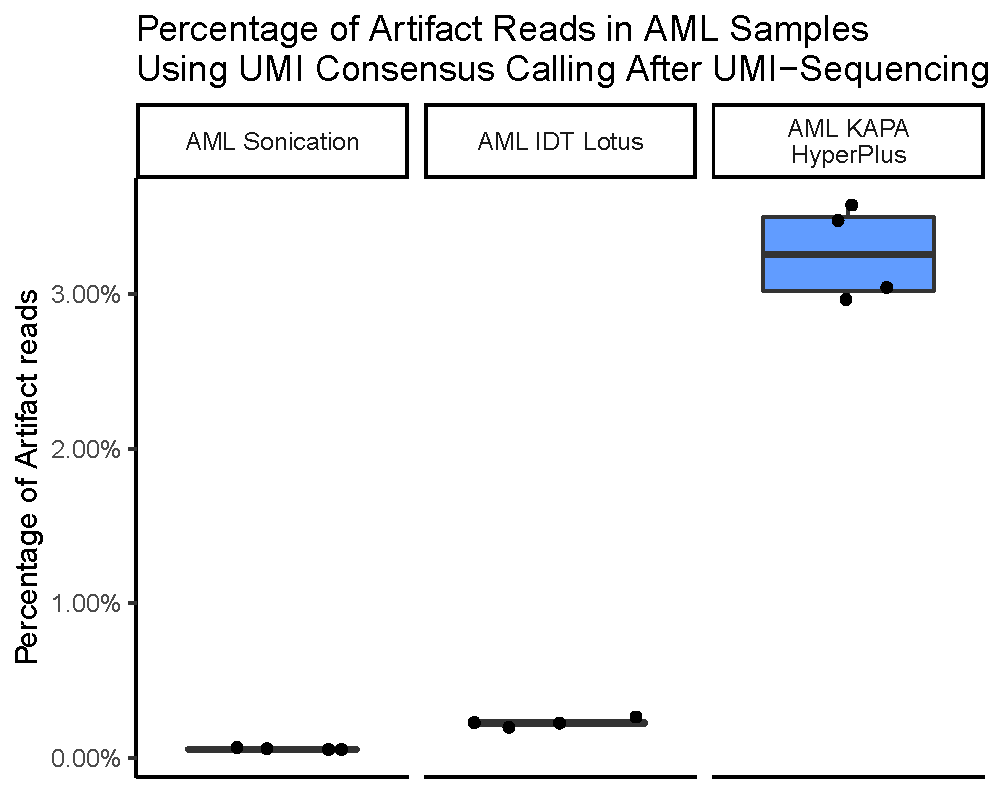

Supp. Figure 1: UMI Consensus calling does not appear to affect overall artifact rates as UMIs are ligated to the source material after fragmentation.
